# Origins and impacts of new mammalian exons

**DOI:** 10.1101/009282

**Authors:** Jason Merkin, Ping Chen, Maria Alexis, Sampsa Hautaniemi, Christopher B. Burge

## Abstract

Mammalian genes are composed of exons, but the evolutionary origins and functions of new internal exons are poorly understood. Here, we analyzed patterns of exon gain using deep cDNA sequencing data from five mammals and one bird, identifying thousands of species- and lineage-specific exons. Most new exons derived from unique rather than repetitive intronic sequence. Unlike exons conserved across mammals, species-specific internal exons were mostly located in 5′ untranslated regions and alternatively spliced. They were associated with upstream intronic deletions, increased nucleosome occupancy, and RNA polymerase II pausing. Genes containing new internal exons had increased gene expression, but only in tissues where the exon was included. Increased expression correlated with level of exon inclusion, promoter proximity, and signatures of cotranscriptional splicing. Together these findings suggest that splicing at the 5′ ends of genes enhances expression and that changes in 5′ end splicing alter gene expression between tissues and between species.

## Introduction

The split structure of eukaryotic genes enables evolutionary changes such as exon gain or loss that are not possible in prokaryotes but may contribute to changes in protein domain structure (Patthy, 2003). The evolutionary gain and loss of introns and exons from genes has been studied in various lineages. Intron gain by unknown mechanisms has been detected in nematodes, fungi and elsewhere (Kiontke et al., 2004, Nielsen et al., 2004), but is rare or nonexistent in mammals, while intron loss has been observed in many lineages (Roy et al., 2003). Similarly, exon loss is detected in various lineages, by mechanisms including genomic deletions and mutational disabling of splice sites (Alekseyenko et al., 2007). Exons have also been gained during evolution by various means including exon duplication (Kondrashov and Koonin, 2001) and acquisition of splice sites and exonic features by an intronic segment (Alekseyenko et al., 2007). The latter process has been most fully characterized for primate-specific exons that have arisen from transposable elements of the Alu family (Lev-Maor et al., 2003, Sorek et al., 2004).

The functional consequences of changes in mRNA splicing patterns can be diverse and have been most widely documented for cases of alternative splicing. For example, inclusion or exclusion of an exon can alter the DNA binding affinity of a transcription factor (Gabut et al., 2011), convert a membrane protein into a soluble protein (Izquierdo et al., 2005), or alter activity and allosteric regulation of an enzyme (Christofk et al., 2008). Evolutionary gain and loss of exons has likely given rise to a similar spectrum of protein functional changes.

Acquisition or loss of an intron can also impact mRNA function. For example, insertion of an intron into a previously intronless expression construct often enhances expression by several fold (or more) in both plant and animal systems in the phenomenon known as intron-mediated enhancement (Callis et al., 1987, Nott et al., 2003), and intronless genes are expressed at lower levels overall (Shabalina et al., 2010). Some effects of introns on mRNA decay (Sureau et al., 2001) or localization (Hachet and Ephrussi, 2004) have been linked to the exon junction complex (EJC), a protein complex deposited just upstream of each exon-exon junction in metazoans that can contribute to mRNA export, translation and stability (Lu and Cullen, 2003, Nott et al., 2003).

Much of what is known about the evolution of mammalian exons has come from the analysis of cDNA fragments known as expressed sequence tags (ESTs). Available EST databases from human and mouse have depths of several million sequences. EST data have certain limitations and biases, including uneven coverage across species (greatest in human and mouse, lower in most others), bias toward cancer in human data, and bias toward brain tissues in mouse, making it difficult to reliably assess splicing in normal tissues or to compare between species (Modrek and Lee, 2003, Roy et al., 2005, Zhang and Chasin, 2006). These issues have contributed to disparities between studies, with reported levels of conservation of alternative splicing between human and mouse ranging from about ¼ to over ¾, depending on the approach that was used (reviewed by (Lareau et al., 2004)), and EST estimates of the fraction of mammalian exons that derive from transposable elements (TEs) range from less than 10% in mouse (Wang et al., 2005) to more than 90% in primates (Zhang and Chasin, 2006).

Subsequent studies increased the number of species and evolutionary distances considered and using phylogenetic information to classify exons by age (Zhang and Chasin, 2006, Alekseyenko et al., 2007). However, splicing patterns in these species generally had to be inferred from multi-species genomic alignments, using presence/absence of AG/GT splice site dinucleotides (which are generally necessary but not sufficient for splicing) to infer splicing/non-splicing of exons, because EST databases were limited or absent from many species. However, using comprehensive RNA-seq data, we observed that presence of AG/GT is not highly predictive of splicing – about half of the exons expressed in mouse but not rat had AG/GT sequences in rat, while the other half did not (Fig. 1). These data imply that exon age classifications based on comparative genomics alone are unreliable, raising questions about conclusions reached using this approach.

**Figure 1.**
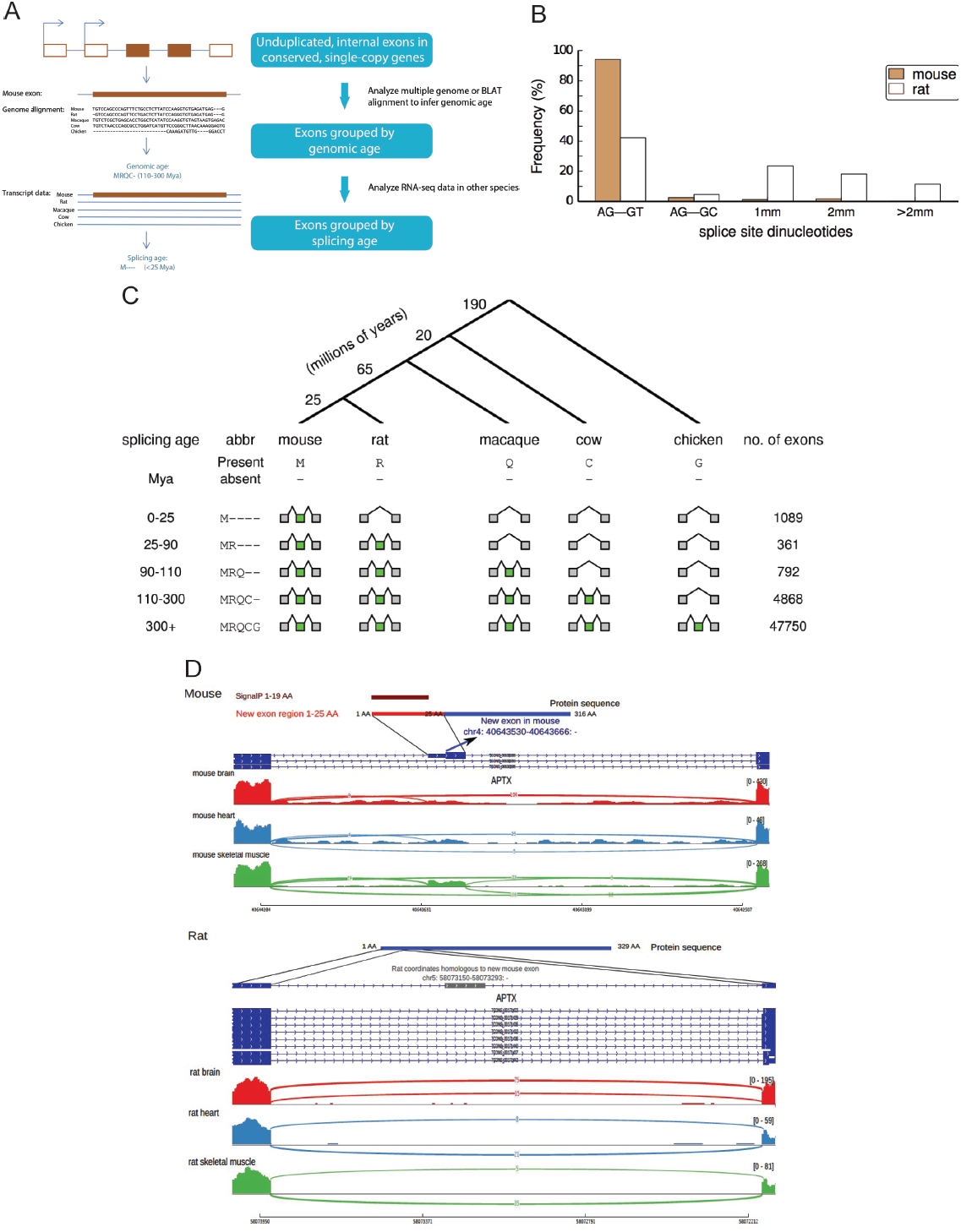
Identification and classification of species- and lineage-specific exons. A. A schematic of our bioinformatic pipeline to identify species- and lineage-specific exons (Methods). We considered all internal exons in each target species (here, mouse) and aligned to other exons in the same gene to exclude cases of exon duplication (Kondrashov and Koonin, 2001). Multiple alignments of orthologous gene sets were used to assign an orthologous region to each exon in other species, and the pattern of genomic presence/absence was used to assign the genomic age by parsimony. Presence/absence of RNA-seq evidence of an overlapping exon in the orthologous region in each species was then used to determine the splicing age, again using parsimony.
B. The proportion of mouse-specific exons and corresponding rat proto-exons with specific dinucleotide sequences at the 3′ and 5′ splice sites (1mm indicates one mismatch relative to the AG-GY consensus, 2mm = 2 mismatches, etc.).
C. Top: a phylogenetic tree presenting the main species used for dating exons and the branch lengths in millions of years. Bottom: mouse exons of increasing splicing age, their pattern of presence/absence in various species, and the number of each class of exons identified.
D. Example of a mouse-specific exon that encodes a predicted N-terminal signal peptide by SignalP (Bendtsen et al., 2004). A portion of the mouse aprataxin (*Aptx*) gene is shown (Ensembl ID ENSMUSG00000028411), together with homologous sequences from rat. Transcript structures shown in dark blue (the gray box in rat is the genomic segment homologous to the mouse-specific exon), with RNA-seq read density from three tissues shown below (arcs represent splice junction reads). See also Figure S1, Table S3.

Only recently has deep RNA-seq analysis been performed systematically across a range of tissues in diverse mammalian species, enabling comprehensive and reliable inference of the phylogenetic distribution of splicing events (Barbosa-Morais et al., 2012, Merkin et al., 2012). In one we performed RNA-seq analysis of 9 diverse organs from 4 mammals and one bird, in biological triplicate (Merkin et al., 2012). The much deeper coverage provided by this RNA-seq dataset – billions of reads totaling hundreds of billions of base pairs (Gbp) for each species, versus millions of sequences totaling at most a few Gbp per species for EST data – and the much more uniform distribution across tissues and species make it much better suited for evolutionary comparisons. The datasets used here and in several previous studies are listed in Supplemental Table S1. In our previous study, we considered exons conserved across mammals and birds, analyzing changes in tissue-specific splicing patterns and conversion between constitutive and alternative splicing. Here, we used these RNA-seq data to identify species- and lineage-specific exons, and assessed their origins, the types of mutations that create them, and their impacts on host genes. Our analyses challenge some previous finding based on EST analyses, uncover an unanticipated role for intronic deletions in exon creation, and provide evidence for a widespread impact of splicing changes on tissue- and species-specific differences in gene expression.

## Results

### Identification of thousands of species- and lineage-specific exons

We combined genomic mappings of our mammalian RNA-seq data with whole-genome alignments to classify exons as species-specific, lineage-specific (e.g., unique to rodents, to primates or to mammals), or ancient (present in both mammals and birds) (Experimental Procedures). These classifications were applied at both the genomic sequence level (“genomic age”) and at the transcript level (“splicing age”) (Fig. 1A), with sequence coverage requirements described in Supplemental Experimental Procedures. Because our primary focus was splicing, we analyzed internal exons – which are regulated by the splicing machinery – rather than first or last exons, which are generally regulated by the transcription or cleavage/polyadenylation machinery, respectively. We also excluded several thousand exons that likely arose from intra-genic duplications, because this class has been thoroughly studied and is fairly well understood (Kondrashov and Koonin, 2001, Gao and Lynch, 2009). Using the principle of parsimony, we assigned both a genomic age and a splicing age to ~60,000 internal exons, restricting our analysis to unduplicated protein-coding genes conserved across these species to facilitate accurate read mapping and assignment of orthology. Here, “genomic age” estimates the duration over which sequences similar to the exon were present in ancestral genomes, while “splicing age” estimates the duration over which these sequences were spliced into mRNAs, based on the RNA-seq data.

Most exons in the analyzed genes predated the split between birds and mammals (~300 million years ago, Mya) in their splicing age (87%) and genomic age (90%). Such exons are designated `MRQCG` using a one-letter code for the five organisms: `M` for present in Mouse, `R` present in Rat, `Q` present in macaQue, `C` present in Cow, `G` present in *Gallus* (chicken) (Fig. 1C). However, we found that creation of novel exons has occurred fairly often during mammalian evolution. For example, we classified 1089 mouse exons as mouse-specific (designated `M----`, with - indicating absence from an organism), as they were detected in RNA-seq data from mouse but not from any other species (Fig. 1C), with 1571 new exons in rat, and 1417 in macaque. Exons in each species, most of which were previously annotated, are listed in Table S2; examples are shown below. Much larger numbers of exons were observed for phylogenetic distributions involving single exon gain and loss events than for those requiring multiple gains/losses (Table S3). The estimated false discovery rate (FDR) for new exons was ~1.5%. This value was estimated using the approach of (Nielsen et al., 2004), in which the rates of gain and loss of a feature (exons here, introns in the previous study) along each branch of a phylogenetic tree are estimated by maximum likelihood from the counts of each phylogenetic pattern of presence/absence, and used to estimate an FDR as the frequency with which parsimony assigns an incorrect age (i.e. an apparent exon gain actually resulted from multiple losses).

Mouse-specific exons were assigned an age of < 25 My, corresponding to the time of divergence between mouse and rat. We also identified ~7000 mouse exons whose splicing was restricted to mammals or to particular mammalian lineages (Fig. 1C). Overall, presence of one or more mouse- or rodent-specific exons was detected in 17% of the ~6300 genes analyzed. An example of a mouse-specific exon that is predicted to encode a signal peptide, and supporting RNA-seq data in mouse and rat, are shown in Figure 1D (see also Fig. S1A). To ask whether species-specific exons occurred at a similar frequency in human, we compared our data to corresponding tissue RNA-seq data from the Illumina Human Body Map 2.0 dataset (Bradley et al., 2012). In the human tissue dataset – which included more tissues, sequenced at lower depth – we identified comparable numbers of exons at each splicing age (Fig. S1B), including 2073 human-specific exons (not observed even in macaque), occurring in 25% of analyzed human genes. Overall, when comparing human genes to their mouse orthologs, we observed that 35% of ortholog pairs differ by presence/absence of one or more exons. The prevalence of such species-specific exons could contribute to functional differences between human and mouse orthologs, complicating extrapolations from mouse models to human phenotypes, though this possibility remains to be explored.

We performed clustering of tissue samples based on tissue-specific splicing patterns of species-specific exons, using the standard “percent spliced in” (PSI) measure, which assesses the fraction of a gene’s mRNAs that include the exon (Experimental Procedures). This analysis revealed robust clustering by tissue of origin across the three mouse strains analyzed (Fig. 2A), with the partial exception of (developmentally related cardiac and skeletal muscle). Many species-specific alternative exons showed predominant inclusion in testis (Fig. 2A). These observations suggest a possible role for germ cell transcription in exon creation, perhaps related to mutations associated with open chromatin and/or transcription-coupled DNA repair (Marques et al., 2005, Levine et al., 2006).

**Figure 2.**
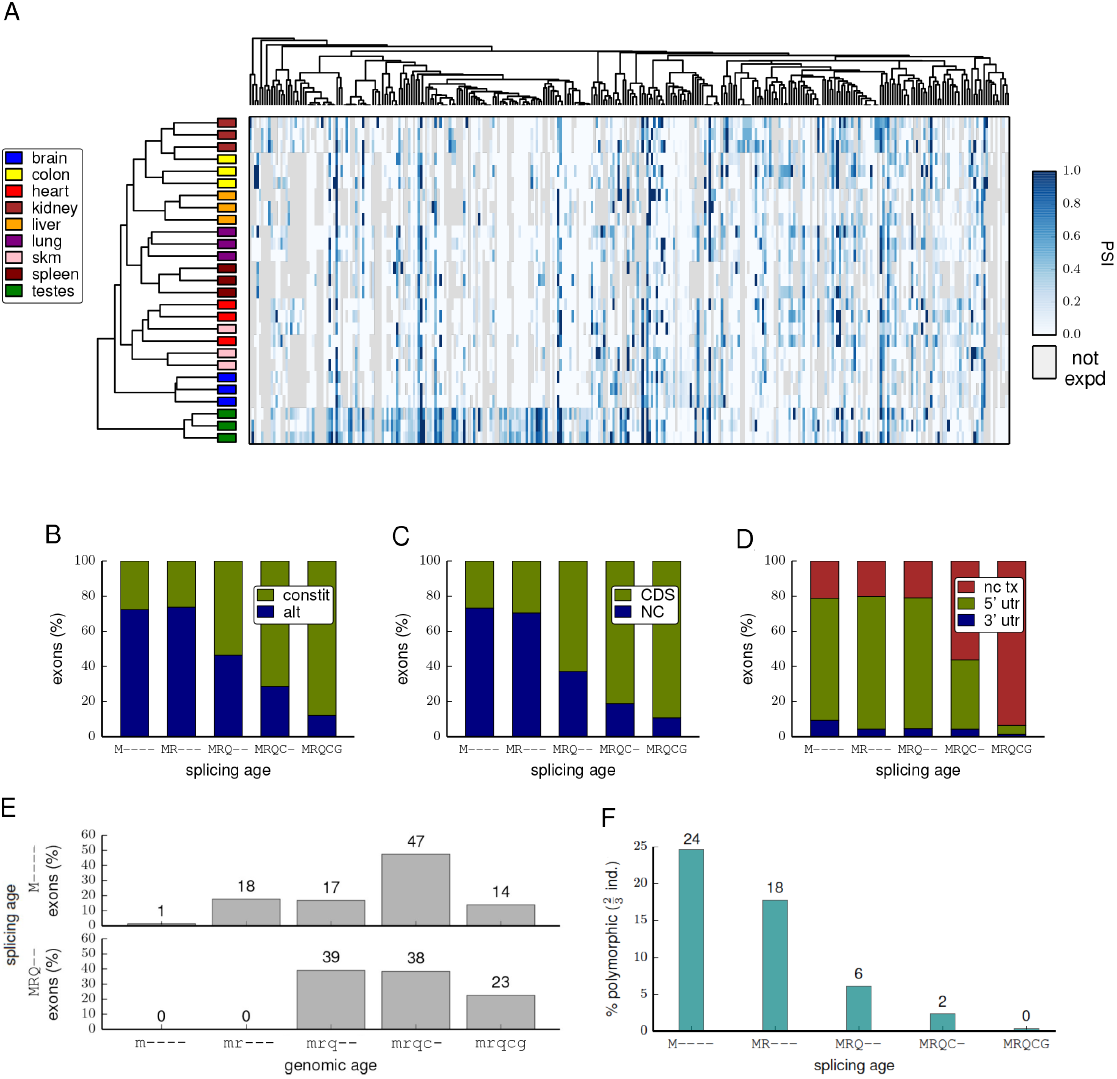
Evolutionarily young exons differ from older exons in many properties. A. Average-linkage hierarchical agglomerative clustering of samples (horizontal axis) or exons (vertical axis) based solely upon PSI values of mouse-specific exons. The tissue of origin of each sample is colored according to the key at left and the PSI value is visualized in the heat map on a white-blue scale (gray indicates gene not expressed in tissue).
B. The proportions of exons of various ages that are alternatively or constitutively spliced.
C. The proportion exons of various ages that contain coding sequence (CDS) or are entirely non-coding (NC) is shown.
D. The proportions of non-coding exons of various ages that are located in non-coding transcripts (nc tx), or in 5′ or 3′ UTRs of coding transcripts.
E. The distributions of genomic ages of exons with splicing ages `M----` or `MRQ--`. Genomic age is represented by the same five-letter code in lowercase.
F. The proportion of mouse exons of various ages that were detected in only 2 out of 3 individuals or where the splicing status (alternative or constitutive) in one individual differed from the other two mice. See also Table S4.

Exons of different evolutionary ages had dramatically different properties. Mouse-specific and other species-specific exons were mostly alternatively spliced, non-coding and located in 5′ UTRs (Fig. 2B-D). Moreover, increasing exon age was consistently associated with constitutive splicing and location within the coding region (Modrek and Lee, 2003, Zhang and Chasin, 2006). No enrichment for interesting categories was observed for the set of genes containing species-specific coding exons (Table S4), suggesting that exon creation occurs with reasonable frequency in diverse types of genes.

To understand the effects of exon inclusion on protein function usually requires extensive experimental analysis. However, functional impact could be inferred in some cases, such as new exons that encode a predicted signal peptide (Fig. 1D) or add a predicted transmembrane domain (Fig. S1A). These two protein motifs have relatively relaxed sequence requirements emphasizing hydrophobicity, suggesting they can evolve quickly.

### Most species-specific exons arose from unique intronic sequences

Considered the genomic ages of recently created exons, approximately 1% of mouse-specific exons arose in sequence detected only in mouse, while 75% of mouse-specific exons arose in sequence that predates the rodent-primate split, despite being spliced exclusively as intron in the other species studied, and the remaining 24% were alignable to rat only (Fig. 2E). Exons with rodent-specific or rodent/primate-specific splicing also could often be aligned to cow or chicken (Fig. 2E and data not shown). Some mouse-specific exons were detected in just two of the three mouse strains analyzed (Fig. 2F).

Notably, we observed that more than 60% of new internal exons in mouse are derived from unique intronic sequence. In most cases, these exons aligned to sequences in the orthologous intron in rat (Fig. 3A). Applying a similar approach to identify new human exons – using criteria designed to allow mapping to repetitive elements (Experimental Procedures) – yielded a similarly high proportion of unique mapping (~54%) (Fig. S2A). Alu elements, a class of primate-specific SINE repeats, have previously been implicated as a major source of new exons in primates (Sorek et al., 2002, Lev-Maor et al., 2003), and an EST-based study concluded that a majority of new human exons are repeat-derived (Zhang and Chasin, 2006). Here, we found that ~19% of exons classified as human-specific overlap with Alus (Fig. S2A). A similar proportion of mouse-specific exons (~18%) overlapped with rodent B elements (Fig. 3B), also above the background genomic frequency of SINEs in this lineage (Fig. 3C and Fig. S2B). Thus, we found that SINEs have contributed to new exon creation to a similar extent in the rodent lineage as in primates (Sela et al., 2007). Many new mouse exons derived from B1 SINE elements aligned to the same positions relative to the antisense B1 consensus (Fig. S2C), consistent with previous studies (Sela et al., 2007).

**Figure 3.**
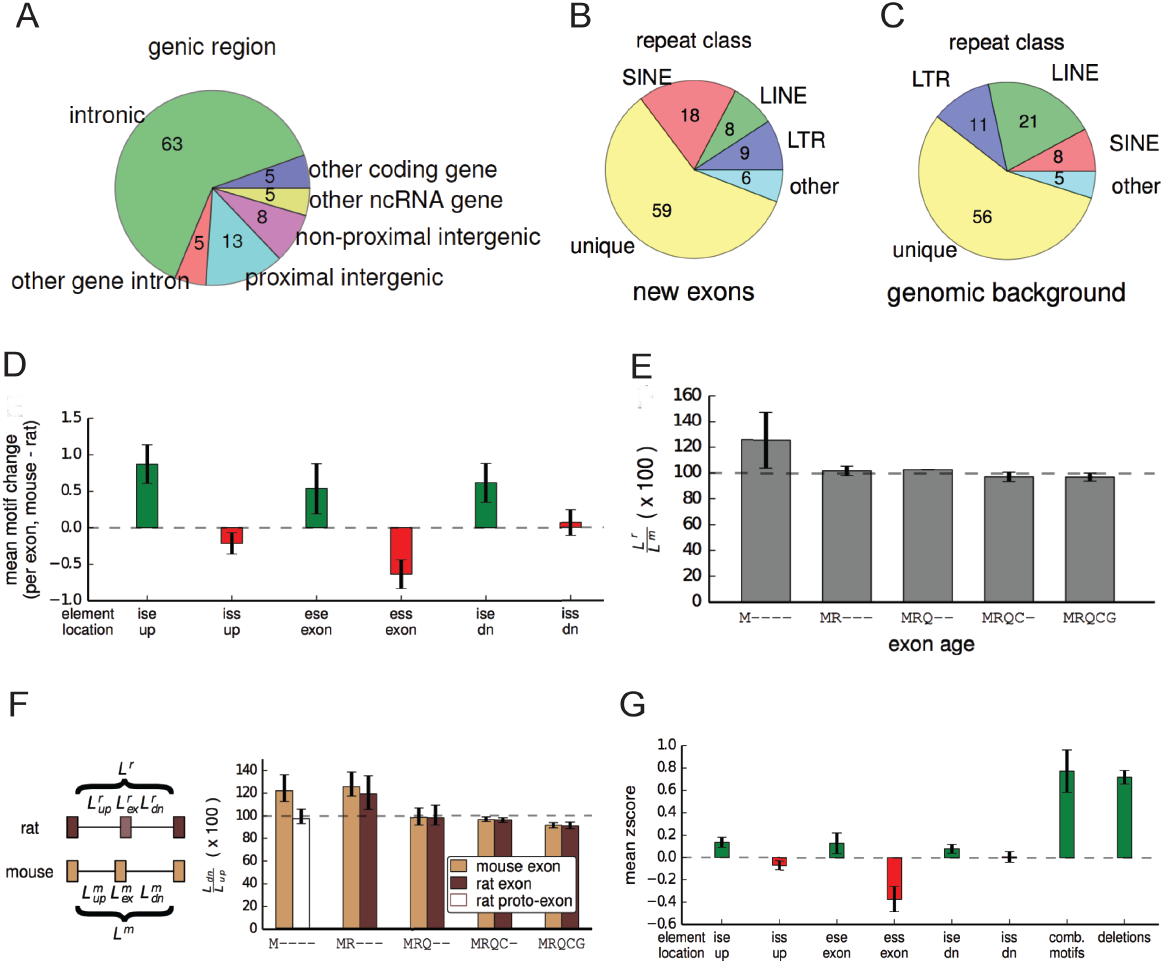
A variety of genomic changes are associated with novel exon splicing. A. Proportion of mouse-specific exons that map to different classes of genomic regions in rat. Mouse-specific exons that mapped to intergenic regions were further classified as proximal intergenic if they were closer to the orthologous gene than any other gene, or otherwise non-proximal intergenic.
B. Proportion of mouse-specific exons that overlap with various classes of repeats.
C. Proportion of mouse genome that belongs to various repeat categories.
D. The change in SRE number in various regions in and around a new exon associated with its creation (mean ± SEM).
E. The change in length of the entire intron region between rat and mouse (see diagram in (F)). The length in rat is plotted as a percentage of the length in mouse (mean ± SEM).
F. The relative length of the downstream intron as a percentage of the upstream intron (mouse) or the downstream aligned intron/region as a percentage of the upstream aligned intron/region (rat) (mean ± SEM). The rat bar in the `M----` class is hatched to indicate that the region is not an exon in rat.
G. The magnitude of each change associated with splicing of `M----` exons was converted into a z-score based upon the distribution of such changes between mouse and rat in `MRQCG` exons. Motifs that are expected to promote splicing are colored in green and changes that are expected to inhibit splicing are shown in red. See also Figures S2, S3, S4.

Our findings indicate that SINEs have contributed 2- to 3-fold fewer new exons than unique genomic sequences in both rodents and primates, contrasting with some previous reports (Zhang and Chasin, 2006). The differences in conclusions presumably result from differences in data sources and methods – e.g., biases in EST coverage of species, tissues and normal versus disease states. Other types of repetitive elements (LINEs, LTRs and others) together have contributed comparable numbers of species-specific exons as SINEs in both human and mouse (Fig. 3B and Fig. S2A).

### Altered splicing motifs and shortened upstream introns are associated with exon creation

Mutations that create or disrupt splice site motifs frequently cause changes in splicing patterns over evolutionary time periods(Lev-Maor et al., 2003, Bradley et al., 2012). Almost all mouse-specific exons contained minimal splice site dinucleotides (AG/ at the 3′ splice site and /GT or /GC at the 5′ splice site) (exceptions: Fig. S2D). However, 47% of homologous “proto-exon” sequences in rat lacked these minimal splicing motifs (Fig. 1B). This observation suggests that mutations that create splice site dinucleotides may contribute to up to about half of exon creation events, but that other types of changes (in *cis* or in *trans*) must explain the remaining cases, where minimal splice site motifs were present in the rat genome but there is no evidence of splicing in rat tissues. Among this subset, strengthening of pre-existing minimal splice site motifs in mouse occurred in about 43% of cases (Fig. S2), but other types of changes must also commonly play a major role in exon creation.

Motifs present in the body of an exon or in the adjacent introns can enhance or suppress exon inclusion (Matlin et al., 2005). We found that mouse-specific exons contain a higher density of exonic splicing enhancer (ESE) motifs and a lower density of exonic splicing silencer (ESS) motifs than their associated rat proto-exons (Fig. 3D). Thus, both gain of enhancing motifs and loss of silencing motifs are likely to contribute to creation and/or maintenance of novel exons. We found an increased density of intronic splicing enhancer (ISE) motifs adjacent to mouse-specific exons relative to homologous rat sequences (but no significant change in intronic silencers), suggesting that changes in flanking sequences may also contribute to exon creation (Fig. 3D). Together, the high frequency of changes to splice site and regulatory motifs associated with new exons suggests that changes in *cis* rather than in *trans* underlie most new exon creation in mammals.

Examining the rates of nucleotide divergence between rat and mouse for mouse exons of different evolutionary ages revealed an interesting pattern (Fig. S3A). While ancient exons exhibited the expected pattern of lower divergence in exons than introns, and even lower divergence in core splice site motifs, mouse-specific exons showed a distinct pattern, in which the central portion of the exon actually diverged more rapidly than flanking intron sequence (which is likely evolving neutrally). This enhanced divergence suggests the existence of positive selection on new exons, perhaps to tune their inclusion levels or optimize their protein-coding or UTR sequence properties.

Intron length is associated with several splicing properties, and longer flanking introns tend to be associated with lower inclusion levels of alternative exons (Yeo et al., 2005). We therefore asked whether species-specific exons were associated with changes in intron length. Notably, we found that the distance between the exons flanking `M----` exons was shorter on average than that flanking homologous rat proto-exons (rat distance exceeded mouse by 1.3-fold on average; interquartile range: 0.9-fold to 1.7-fold; Fig. 3E). The distance between the exons flanking `-R---` exons was even more biased, with exons flanking mouse proto-exons longer by 1.7-fold on average (Fig. S3B). These observations suggest that substantial changes in intron length often accompany exon creation. Comparing the lengths of the upstream and downstream introns flanking species-specific exons, we observed that the intron downstream of `M----` exons was 1.2-fold longer on average than the upstream intron, compared to no difference between the regions flanking rat proto-exons (Fig. 3F), with a somewhat smaller effect observed for `-R---` exons (Fig. S3C). In rodent-specific exons, a similar bias towards presence of a longer downstream intron was observed in both mouse and rat. Comparison to an outgroup (macaque) indicated that the differences in flanking intron length most often reflect upstream deletions rather than downstream insertions in the rodent that acquired a new exon (Fig. S3D, S3E). Older groups of exons showed no such bias, suggesting that exons may acquire tolerance for expansion of the upstream intron over time as other splicing determinants strengthen. Together, these data argue that deletions upstream of proto-exons favor creation and/or maintenance of novel exons. Previously, shortening of an upstream intron was associated with enhancement of exon inclusion in a minigene reporter context (Fox-Walsh et al., 2005), perhaps by enhancing intron definition or exon juxtaposition following exon definition, but the generality of this effect and its evolutionary impact have not been explored.

To compare the relative magnitudes of genomic changes associated with species-specific exons, we converted them all to z-scores, scaled by the standard deviation of the differences observed between homologous mouse and rat ancient (`MRQCG`) exons. Each sequence motif type had a relatively small z-score (< 0.4). However, upstream intronic deletions had an average z-score of 0.75, comparable to the sum of the z-scores of all sequence motifs analyzed (Fig. 3G). This observation suggests that upstream intronic deletions may contribute to exon creation to a similar extent as changes in known classes of splicing regulatory elements (SREs; see also Fig. S4 and Supp. Experimental Procedures). New exons associated with upstream intronic deletions had a similar distribution of splice site dinucleotides as other new exons but lack the increased ESE frequency observed for other new exons (Fig. S4). This observation is consistent with a model in which upstream intron shortening promotes exon inclusion to such an extent that selective pressure to create or preserve ESEs is relaxed.

### Upstream indels are associated with nucleosome occupancy and RNA Pol II pausing over novel exons

Previous studies have suggested functional links between nucleosomes and splicing. Nucleosomes are often positioned near the centers of internal exons, exon-associated nucleosomes have higher density of the H3K36me3 histone modification (Schwartz et al., 2009, Spies et al., 2009), and histone modifications can impact recruitment of splicing factors (Luco et al., 2010). Conceivably, insertions/deletions (indels) in flanking introns such as those observed in Figure 3E,F might impact splicing by altering nucleosome positioning near an exon. To explore this possibility, we used published micrococcal nuclease- (MNase-) sequencing data from digestion of chromatin to identify nucleosome-protected regions in the vicinity of mouse-specific exons. We observed a stronger enrichment for nucleosome positioning over those mouse-specific exons which had shortened upstream introns (relative to rat) compared to ancient exons, or to mouse-specific exons without upstream shortening, or to mouse proto-exons orthologous to rat-specific exons (Fig. 4A; P < 10^−4^ for all three comparisons by Kolmogorov-Smirnov (KS) test). This association suggested a connection between upstream deletions and changes in nucleosome positions. While indels in either the upstream or downstream intron could potentially impact nucleosome positioning on an exon, upstream deletions may be more likely than other types of indels to promote exon inclusion because a shorter intron promotes intron definition or exon juxtaposition, possibly augmented by effects on exon-proximal nucleosome positioning.

**Figure 4.**
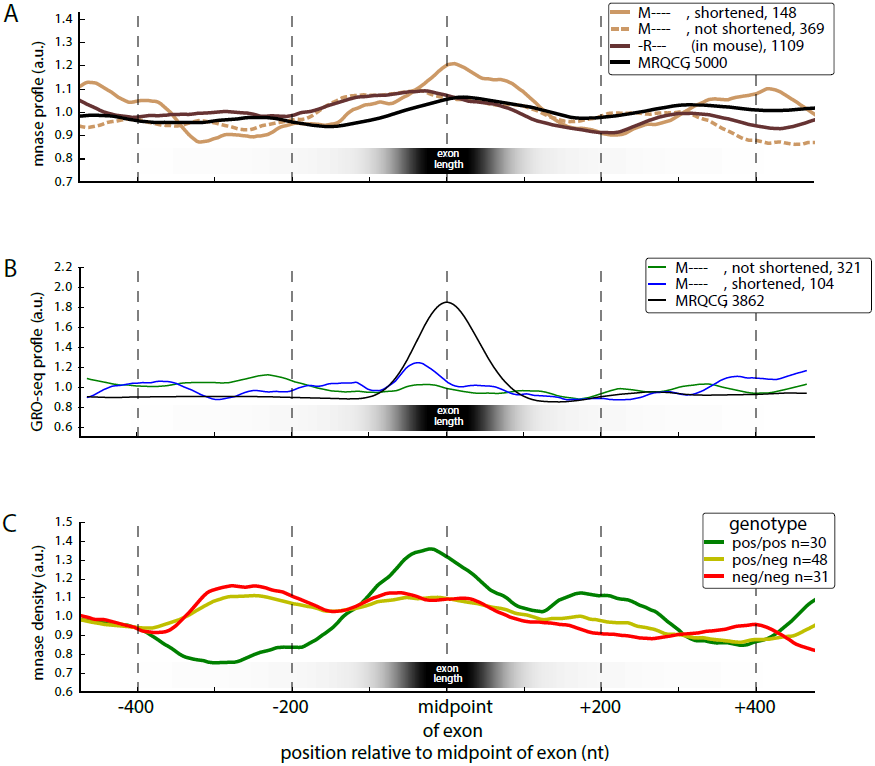
Upstream intronic deletions are associated with increased exonic nucleosome occupancy and transcription pausing. A. Nucleosome positioning (measured by MNase protection) around various sets of exons.
B. Density of global run-on sequencing (GRO-seq) reads, showing the position of elongating RNA Pol II.
C. Nucleosome positioning (measured by protection from MNase treatment) around exons with a structural sQTL in the upstream intron binned by sQTL genotype. See also Figure S4, S5.

We hypothesized that changes promoting stronger nucleosome positioning on novel exons might slow polymerase elongation (Bentley, 2014) and thereby act to promote splicing (Gunderson et al., 2011, Zhou et al., 2011, Ip et al., 2011). To test this hypothesis, we used global run-on-sequencing (GRO-seq) data, which detects nascent transcription, from a recent study (Kaikkonen et al., 2013). We observed a strong GRO-seq peak over ancient exons, almost twice the background level (Fig. 4B; P < 10^−4^, KS test), suggesting that polymerases decelerate by roughly twofold while transcribing through these exons; much smaller effects in this direction have been observed using Pol II ChIP-seq data (Schwartz et al., 2009, Spies et al., 2009). When considering mouse-specific exons with an upstream intronic deletion, we observed a GRO-seq peak ~37% above the nearly flat background observed in mouse-specific exons without upstream deletions (P < 10^−4^, KS test). Thus, although confirmation is needed, the observations above suggest a model in which upstream deletions enhance inclusion of new exons in part by promoting nucleosome positioning and slowing polymerase elongation.

Recent studies of the genetic basis for gene expression variation have also identified thousands of genetic variants associated with altered levels of splicing (splicing-quantitative trait loci or sQTLs) between human individuals (Pickrell et al., 2010, Lalonde et al., 2011, Lappalainen et al., 2013). To explore connections between indels, nucleosomes and splicing, we used sQTLs identified in genotyped human lymphoblastoid cell lines studied by the GEUVADIS Consortium (Lappalainen et al., 2013) and MNase-seq data from a subset of these individuals (Gaffney et al., 2012) to assess nucleosome positioning in individuals with different sQTL genotypes. For upstream indel sQTLs we observed that the genotype associated with increased exon inclusion also had increased nucleosome density in the vicinity of the associated exon (Fig. 4C; P < 0.01, KS test). We also found that stronger differences in nucleosome density in the vicinity of the new exon were associated with indel sQTLs located closer to the affected exon, as expected if these indels directly impact nucleosome placement near exons (Fig. S5). Together, the data in Figure 4 implicate upstream intronic indels in alterations to nucleosome positioning and splicing between individuals and species.

### Splicing of species-specific exons is associated with increased gene expression

We next asked what effects new exons have on the genes in which they arise. Since the majority of species-specific exons we identified were non-coding (Fig. 2C), we examined effects on gene expression. Intron-mediated enhancement is a well-established though incompletely understood phenomenon in which introduction of a (possibly heterologous) intron into an intronless gene or minigene often leads to higher expression of the gene (Mascarenhas et al., 1990, Jonsson et al., 1992), through effects on mRNA export, cleavage/polyadenylation, message stability or other mRNA properties (Nott et al., 2003, Lu and Cullen, 2003). Here we studied a different situation, in which an already multi-exonic gene acquires a new internal exon within an intron, incrementing the count of introns and exons in the transcript by one. We observed significantly higher expression of genes containing mouse-specific exons in mouse tissues relative to their rat orthologs in corresponding rat tissues (Fig. 5A). This effect was specific to those mouse tissues where the new exon was included (i.e. spliced in to the mRNA), consistent with a positive effect of splicing on steady state expression levels (Fig. 5A). The inclusion of a new exon was associated with an increase in gene expression of 10% overall (Fig. 5A, inset), with much larger effects observed for specific subsets of new exons (see below). The distribution of splice junction read data supported direct effects of splicing, ruling out alternative explanations involving new promoters (Fig. S6A).

**Figure 5.**
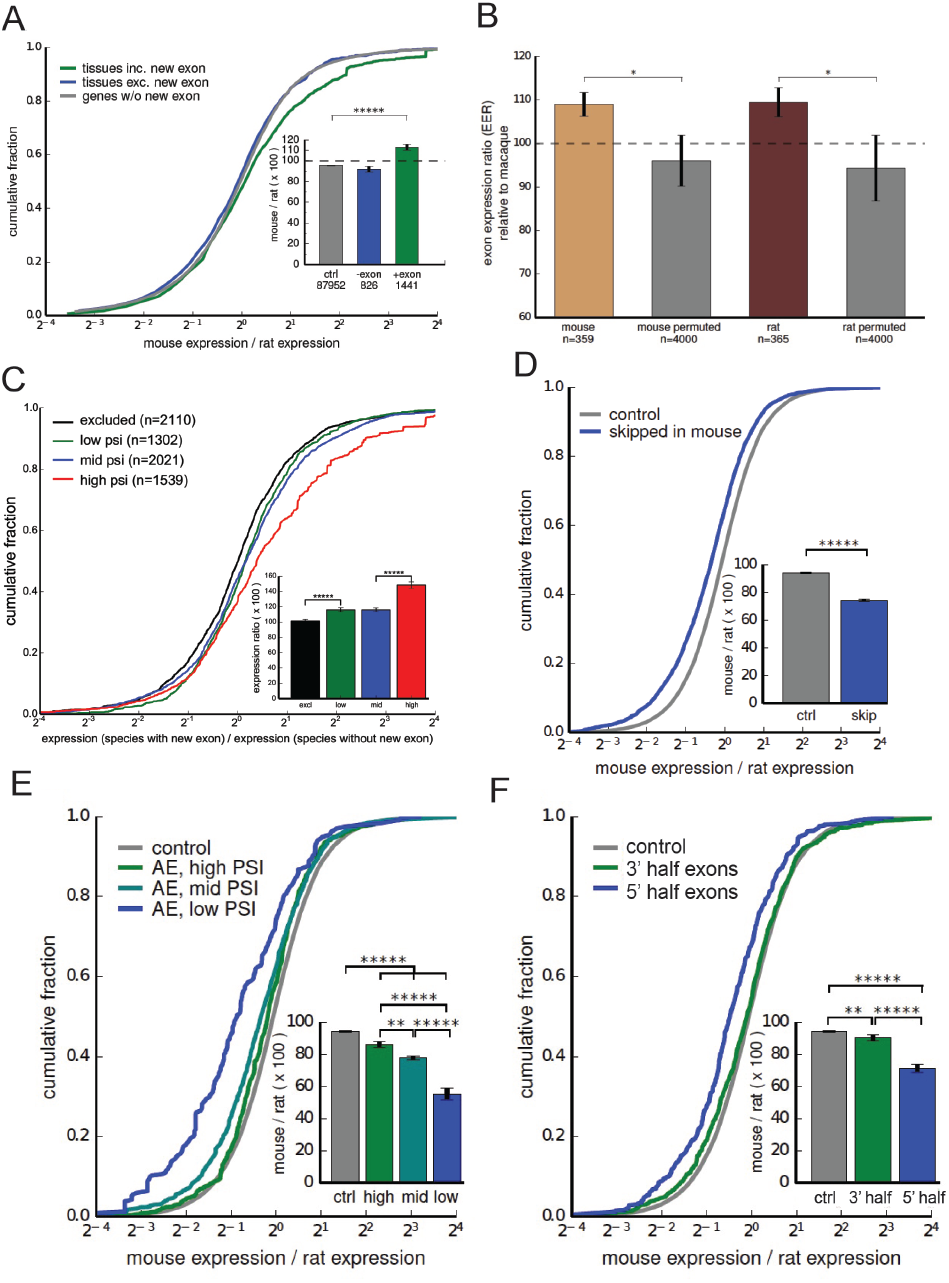
Inclusion of new exons is associated with increased species-specific gene expression changes. Throughout the figure, statistical significance by Mann-Whitney U test is indicated by asterisks (* P < 0.05, ** P < 0.01, *** P < 0.001, **** P < 0.0001, ***** P < 0.00001). A. Fold change in gene expression between mouse and rat. Inset: mean ± SEM of displayed distributions.
B. Mean EEI values, calculated as the ratio of the mean gene expression in tissues where a novel exon is included to the mean expression in tissues where inclusion of the exon is not detected. This ratio is calculated in a related species with matched tissues, and the ratio of these two values is plotted (mean ± SEM).
C. Fold change in gene expression of mouse- and rat-specific exons between species with new exons (mouse, rat) and aligned closest species without new exon (rat, mouse, respectively), binned by the PSI of the exon in the tissue.
D. Fold change in gene expression between mouse and rat in genes where an ancestrally present exon has become skipped in mouse.
E. Fold change in gene expression between mouse and rat in genes where an old exon has become skipped in mouse, binned by the PSI of the exon in the tissue.
F. Fold change in gene expression between mouse and rat in genes where an old exon has become skipped in mouse, binned by location of the exon within the gene. See also Figure S6.

The association between splicing and gene expression can also be assessed by the ratio of the mean expression in tissues where the exon is included to the mean expression in tissues where the exon is excluded, a measure we call the “exon-associated expression index” (EEI). Dividing the EEI in the species that contains the exon to the ratio of mean expression values in the same pairs of tissues in the species lacking the exon yields an “exon-associated expression ratio” (EER), with the log of the EER distributed symmetrically around zero under the null hypothesis that splicing of the new exon does not affect gene expression. This approach controls for various technical factors that could impact comparison of expression levels between different species. Comparing EER values for genes containing mouse-specific exons or rat-specific exons to controls (Fig. 5B), we observed significantly elevated ratios (about 1.1) in both cases, consistent with the 10% increase in expression observed using the simpler approach of Figure 5A, providing support for the alternative hypothesis that splicing in of new exons enhances gene expression.

If splicing in of new exons directly enhances expression then higher PSI values should be associated with greater increase in expression. Consistently, we observed substantial increases in expression (averaging ~50%) for new exons with the highest PSI values in mouse, and also independently in rat (combined data shown in Fig. 5C), supporting the model in which splicing in of new exons directly enhances expression. To further explore this phenomenon, we considered another set of exons: exons whose presence in the transcriptome is ancient, but which undergo exon skipping only in mouse (Merkin et al., 2012). We observed that genes containing such exons had lower gene expression in mouse than their rat orthologs, consistent reduced splicing contributing to reduced expression (Fig. 5D). Furthermore, we observed a dose-dependent effect, in which reduced exon inclusion (lower PSI values) were associated with a stronger decrease in gene expression (Fig. 5E). We also observed a positional effect, in which the strongest expression differences were associated with exons located in the 5′ end of genes, with little effect observed for exons located at gene 3′ ends (Fig. 5F). Most of the species-specific exons analyzed in Figure 5A were located near the 5′ ends of genes (Fig. 2D). A certain minimum level of expression is required to detect exon skipping and to detect novel exons. However, we took measures to counteract such biases (Experimental Procedures), and we note that any such bias would tend to oppose the effect on expression observed in Figure 5D-F.

Splicing of pre-mRNAs can impact cytoplasmic mRNA decay (e.g., mediated by effects of EJC proteins deposited after splicing), but recent studies have also pointed to connections between splicing and transcription in various systems. These connections include reports that splicing can influence transcription initiation (Damgaard et al., 2008) or elongation (Singh and Padgett, 2009, Lin et al., 2008), and recent evidence in yeast of a checkpoint in which the polymerase pauses during intron transcription until partial spliceosome assembly occurs (Chathoth et al., 2014). A direct effect of splicing on transcription would be expected to occur only for exons and introns that are cotranscriptionally spliced, while an effect on cytoplasmic mRNA decay should be independent of co-versus post-transcriptional splicing. To address this issue we used a measure of post-transcriptional splicing, here termed the incomplete splicing ratio (ISR) (Brugiolo et al., 2013) (Methods). ISR measures the ratio of intronic to exonic transcript reads, with higher values indicating more post-transcriptional splicing and lower values more co-transcriptional splicing; the distribution of ISR values is shown in Figure 6A. Notably, we observed that the decrease in gene expression in mouse relative to rat associated with mouse-specific exon skipping was greatest for exons with low ISR, and nonexistent for exons with high ISR, with moderate decrease observed for the exons with middle levels of ISR (Fig. 6B). In other words, the association between reduced splicing and reduced expression was correlated with the extent of cotranscriptional splicing. This observation makes sense if there is direct linkage between splicing and transcription but is difficult to reconcile with models involving effects of splicing on cytoplasmic decay. Interestingly, cotranscriptional splicing is reported to be much more efficient for introns located far from 3′ ends of genes (Khodor et al., 2012). Therefore, the positive association between expression and splicing at the 5′ but not the 3′ end of genes observed above (Fig. 5F) may be related to the extent of cotranscriptional splicing.

**Figure 6.**
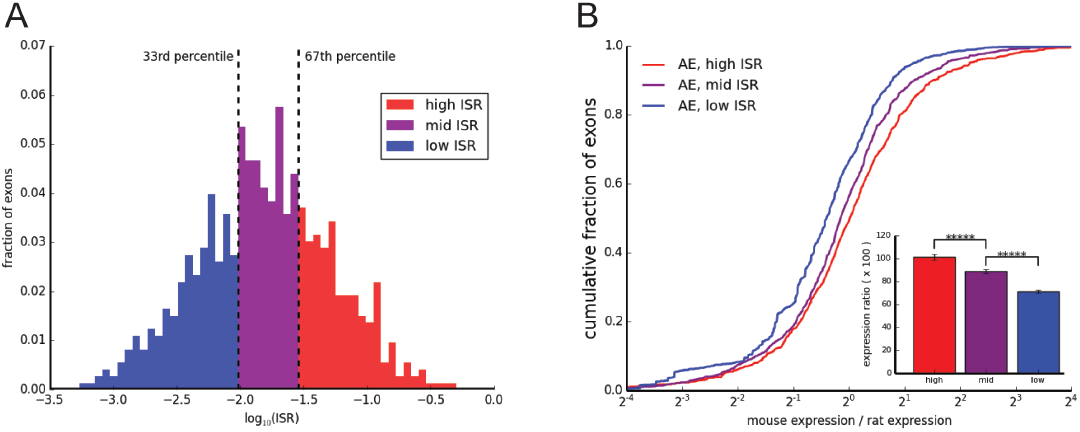
Increase in species-specific gene expression is associated with a lower ISR (Incomplete Splicing Ratio). A. Histogram of ISR values (on a log-scale) of rat constitutive exons which are ancestrally present but have become skipped in mouse.
B. Fold change in gene expression between mouse and rat in genes where an ancient exon has become skipped in mouse, binned by ISR as in (A). Statistical significance by Mann-Whitney U test is indicated by asterisks, as in Figure 5 (***** P < 0.00001).

## Discussion

Here, we identified and analyzed thousands of orthologous gene sets differing by evolutionary gain of a single exon, most often alternatively spliced and located in the 5′ UTR, offering a natural system in which to examine the effects of exons and splicing on host genes. Various factors may contribute to the bias for new exons to occur in 5′ UTRs, including the greater length of first introns relative to later introns (Kriventseva and Gelfand, 1999), the low frequency of 3′ UTR introns (Giorgi et al., 2007), and the potential for some new coding and 3′ UTR exons to destabilize messages by eliciting NMD. Several phenomena reported here, including the strong association between upstream deletions and emergence of new exons (Fig. 3), and the associated changes in chromatin structure and polymerase dynamics (Fig. 4) have not been observed in previous EST-based studies, and our conclusion that most new exons derive from unique sequences (Fig. 2) differs from that obtained in some previous EST-based studies (Zhang and Chasin, 2006), emphasizing the difference that use of comprehensive RNA-seq data can make.

Most intriguing was the observation of a consistent association between increased splicing at the 5′ ends of transcripts and increased mRNA expression, a finding that has important implications. Analyzing genes that exhibit species-specific increases in gene expression, we observed substantial enrichment (1.9-fold in mouse, 1.3-fold in rat) for genes containing species-specific exons (Fig. S6B). This observation, together with the results of Figure 5, suggest that splicing of 5′ end exons plays a general role in promoting gene expression and that gain and loss of 5′ UTR exons and changes in their splicing patterns contribute substantively to evolutionary and tissue-specific differences in gene expression. The observed relationship between exon gain and expression increase is consistent with the general trend observed previously in which more ancient human genes have both higher intron density and higher expression (Wolf et al., 2009).

Our approach of globally analyzing changes in gene structure and expression across thousands of genes in several species and tissues is complementary to more targeted molecular approaches aimed at dissecting the relationship between exon structure/splicing and gene expression. Averaging over hundreds or thousands of genes enables detection of general trends above the background of idiosyncratic features of individual transcripts that can confound targeted mutational analysis, though the discernment of cause and effect is more challenging.

Steady-state gene expression is determined by the balance between the rate of production of mRNAs in the nucleus and the rate of decay, occurring predominantly in the cytoplasm. Consistently, studies of mRNA stability have observed a strong correlation between the density of exon junctions in the open reading frame and mRNA half-life (Sharova et al., 2009, Spies et al., 2013). Despite the initial plausibility of mRNA decay effects, our finding that it is predominantly exons with signatures of cotranscriptional splicing whose splicing is associated with increased gene expression (Fig. 6) argue for a different model, instead suggesting a direct positive link between splicing and transcription in mammalian nuclei. This link – which will require direct confirmation – might reflect the effects of known or unknown mechanisms. For example, U1 snRNP exerts inhibitory effects on premature cleavage and polyadenylation, in a phenomenon known as “telescripting” (Berg et al., 2012), which might well be enhanced by recruitment of U1 snRNP to a novel exon. Alternatively, SR proteins might link splicing to transcription, as these proteins have been implicated in control of transcription elongation as well as splicing (Ji et al., 2013). Independent of mechanism, artificial enhancement of 5′ UTR splicing – via antisense oligonucleotides or other means – might offer a viable means to enhance gene expression for research or therapeutic purposes.

## Methods

### RNA-seq and genome builds

Data from mouse, rat, rhesus, cow, and chicken were mapped to mm9, rn4, rhemac2, bostau4, and galgal3 respectively, and processed as in (Merkin et al., 2012) using TopHat (Trapnell et al., 2009) and Cufflinks (Trapnell et al., 2012).

### Assignment of ages to exons

We used exons from (Merkin et al., 2012), where exons were defined as having FPKM ≥ 2 and meeting splice site junction read requirements implicit in the TopHat mapping. Alignment of exons to other species were collected using whole genome alignments generated by PECAN and EPO (Paten et al., 2008) and pairwise alignments from BLASTZ (Schwartz et al., 2003). Genomic and splicing ages were defined by the pattern of species with genomic regions aligned to the exon or with an expressed region in the orthologous gene overlapping the aligned region, respectively. We used parsimony to find the minimum number of changes that can explain these patterns (Alekseyenko et al., 2007, Roy et al., 2008).

### Basic exon properties

Exons with 0% < PSI < 97% in at least 1 tissue and 2 individuals were categorized as skipped exons (SE). Exons with PSI > 97% in all expressed tissues were defined as constitutive exons (CE), if the gene was expressed in at least 3 tissues and 2 individuals. ORFs (open reading frames) were annotated as in (Merkin et al., 2012) and used to classify exons as coding, 5′ UTR, 3′ UTR, and non-coding.

### Genomic sources of new exons

Exons were categorized as intronic, proximal intergenic, non-proximal intergenic, other coding gene, other intron and other ncRNA gene if their aligned region in the closest species was located in the intronic regions of the same gene, intergenic regions but closer to the orthologous gene than any other gene, other intergenic regions, exonic regions of other genes, intronic regions of other genes and other regions of ncRNA, respectively. Using RepeatMasker, exons were categorized as containing SINEs, LINEs, LTRs, other repeats (rarer categories), or designated as “unique” if not overlapping any repeat class.

### Splice site and SRE analysis

The dinucleotide frequencies of the 5′ and 3′ splice sites of mouse new exons and their aligned regions in rat were compared in Figure 1D. In Figure 3D, the 100 nt up and downstream of each mouse exon (or aligned region in rat) was considered for searching for intronic SREs. The entire exon was searched for exonic SREs.

### Intron length analyses

For each exon age, the sum of lengths of each mouse exon and its flanking introns were compared to the corresponding sum for the rat exon or proto-exon (Fig. 3E). In Figure 3F, the length of the downstream mouse intron was divided by the length of the upstream mouse intron (the same was done for the rat exon or proto-exon).

### Z-score conversion for comparisons

We determined the empirical distribution of the changes considered (changes in intronic or exonic splicing enhancers or silencers, or deletions) in the ancient exons (MRQCG). We then calculated a z-score for each change for each new exon using the empirical mean and standard deviation for ancient exons (Fig. 3G).

### Nucleosome localization and GRO-seq analyses

We used MNase-seq data from (Gaffney et al., 2012) and GRO-seq data from (Kaikkonen et al., 2013) and determined the read density in a 1kb window for each exon. These profiles were normalized, average, smoothed, and centered on the exon midpoint. To investigate the impact of intronic structural variants on nucleosome localization (Fig. 4C), we used the MNase-seq data above and corresponding sQTL (Lappalainen et al., 2013) and genotype data (Abecasis et al., 2012). We then compiled the MNase profiles of individuals with genotypes representing shorter and longer upstream introns.

### New exon inclusion and species-specific expression changes

Gene expression in mouse was compared to that in rat by taking the ratio of mouse to rat expression. In Figure 5A, we considered genes with a new exon, grouped by whether the exon was included or not in a given tissue. In Figure 5C, we considered mouse- and rat-specific exons, taking the expression ratio relative to rat and mouse respectively. We also compared the expression ratio for ancient exons included in rat but skipped in mouse (Fig. 5D), grouped by PSI (Fig. 5E) or position within the gene (Fig. 5E).

The intra-species expression ratio is calculated by dividing the mean mouse gene expression in tissues where the exon is included by the mean expression in the other tissues. This ratio was then calculated in rat, matching the tissues in the fore- and background, and the ratio of these two values was analyzed. This statistic was recalculated with shuffled tissue labels as a control.

The incomplete splicing ratio (ISR) was determined for the exons used in Figure 5D. The ISR of an exon is calculated by dividing the mean read density in its flanking introns by that of its neighboring exons, averaged across tissues.

## Author Contributions

Initial conception of project (J. M., P. C., S. H. and C. B. B.), analyses related to Figures 1-3 (P. C. and J. M.), analyses related to Figure 4 (J. M.), analyses related to Figure 5 (J. M. and M. A.), analyses related to Figure 6 and Table S1 (M. A.), analyses related to other supplemental material (P. C. and J. M.), interpretation of data (all authors), writing of manuscript (all authors), management of project (C. B. B.).

## References

Abecasis, G. R., Auton, A., Brooks, L. D., Depristo, M. A., Durbin, R. M., Handsaker, R. E., Kang, H. M., Marth, G. T. & McVean, G. A. 2012. An integrated map of genetic variation from 1,092 human genomes. Nature, 491, 56–65.

Alekseyenko, A. V., Kim, N. & Lee, C. J. 2007. Global analysis of exon creation versus loss and the role of alternative splicing in 17 vertebrate genomes. RNA, 13, 661–70.

Barbosa-Morais, N. L., Irimia, M., Pan, Q., Xiong, H. Y., Gueroussov, S., Lee, L. J., Slobodeniuc, V., Kutter, C., Watt, S., Colak, R., Kim, T., Misquitta-Ali, C. M., Wilson, M. D., Kim, P. M., Odom, D. T., Frey, B. J. & Blencowe, B. J. 2012. The evolutionary landscape of alternative splicing in vertebrate species. Science, 338, 1587–93.

Bendtsen, J. D., Nielsen, H., Von Heijne, G. & Brunak, S. 2004. Improved prediction of signal peptides: SignalP 3.0. Journal of molecular biology, 340, 783–95.

Bentley, D. L. 2014. Coupling mRNA processing with transcription in time and space. Nature reviews. Genetics, 15, 163–75.

Berg, M. G., Singh, L. N., Younis, I., Liu, Q., Pinto, A. M., Kaida, D., Zhang, Z., Cho, S., Sherrill-Mix, S., Wan, L. & Dreyfuss, G. 2012. U1 snRNP determines mRNA length and regulates isoform expression. Cell, 150, 53–64.

Bradley, R. K., Merkin, J., Lambert, N. J. & Burge, C. B. 2012. Alternative splicing of RNA triplets is often regulated and accelerates proteome evolution. PLoS biology, 10, e1001229.

Brugiolo, M., Herzel, L. & Neugebauer, K. M. 2013. Counting on co-transcriptional splicing. F1000Prime Rep, 5, 9.

Callis, J., Fromm, M. & Walbot, V. 1987. Introns increase gene expression in cultured maize cells. Genes Dev, 1, 1183–200.

Chathoth, K. T., Barrass, J. D., Webb, S. & Beggs, J. D. 2014. A splicing-dependent transcriptional checkpoint associated with prespliceosome formation. Mol Cell, 53, 779–90.

Christofk, H. R., Vander Heiden, M. G., Wu, N., Asara, J. M. & Cantley, L. C. 2008. Pyruvate kinase M2 is a phosphotyrosine-binding protein. Nature, 452, 181–6.

Damgaard, C. K., Kahns, S., Lykke-Andersen, S., Nielsen, A. L., Jensen, T. H. & Kjems, J. 2008. A 5′ splice site enhances the recruitment of basal transcription initiation factors in vivo. Mol Cell, 29, 271–8.

Fox-Walsh, K. L., Dou, Y., Lam, B. J., Hung, S. P., Baldi, P. F. & Hertel, K. J. 2005. The architecture of pre-mRNAs affects mechanisms of splice-site pairing. Proc Natl Acad Sci U S A, 102, 16176–81.

Gabut, M., Samavarchi-Tehrani, P., Wang, X., Slobodeniuc, V., O’Hanlon, D., Sung, H. K., Alvarez, M., Talukder, S., Pan, Q., Mazzoni, E. O., Nedelec, S., Wichterle, H., Woltjen, K., Hughes, T. R., Zandstra, P. W., Nagy, A., Wrana, J. L. & Blencowe, B. J. 2011. An alternative splicing switch regulates embryonic stem cell pluripotency and reprogramming. Cell, 147, 132–46.

Gaffney, D. J., McVicker, G., Pai, A. A., Fondufe-Mittendorf, Y. N., Lewellen, N., Michelini, K., Widom, J., Gilad, Y. & Pritchard, J. K. 2012. Controls of nucleosome positioning in the human genome. PLoS genetics, 8, e1003036.

Gao, X. & Lynch, M. 2009. Ubiquitous internal gene duplication and intron creation in eukaryotes. Proc Natl Acad Sci U S A, 106, 20818–23.

Giorgi, C., Yeo, G. W., Stone, M. E., Katz, D. B., Burge, C., Turrigiano, G. & Moore, M. J. 2007. The EJC factor eIF4AIII modulates synaptic strength and neuronal protein expression. Cell, 130, 179–91.

Gunderson, F. Q., Merkhofer, E. C. & Johnson, T. L. 2011. Dynamic histone acetylation is critical for cotranscriptional spliceosome assembly and spliceosomal rearrangements. Proc Natl Acad Sci U S A, 108, 2004–9.

Hachet, O. & Ephrussi, A. 2004. Splicing of oskar RNA in the nucleus is coupled to its cytoplasmic localization. Nature, 428, 959–63.

Ip, J. Y., Schmidt, D., Pan, Q., Ramani, A. K., Fraser, A. G., Odom, D. T. & Blencowe, B. J. 2011. Global impact of RNA polymerase II elongation inhibition on alternative splicing regulation. Genome research, 21, 390–401.

Izquierdo, J. M., Majos, N., Bonnal, S., Martinez, C., Castelo, R., Guigo, R., Bilbao, D. & Valcarcel, J. 2005. Regulation of Fas alternative splicing by antagonistic effects of TIA-1 and PTB on exon definition. Mol Cell, 19, 475–84.

Ji, X., Zhou, Y., Pandit, S., Huang, J., Li, H., Lin, C. Y., Xiao, R., Burge, C. B. & Fu, X. D. 2013. SR proteins collaborate with 7SK and promoter-associated nascent RNA to release paused polymerase. Cell, 153, 855–68.

Jonsson, J. J., Foresman, M. D., Wilson, N. & McIvor, R. S. 1992. Intron requirement for expression of the human purine nucleoside phosphorylase gene. Nucleic acids research, 20, 3191–8.

Kaikkonen, M. U., Spann, N. J., Heinz, S., Romanoski, C. E., Allison, K. A., Stender, J. D., Chun, H. B., Tough, D. F., Prinjha, R. K., Benner, C. & Glass, C. K. 2013. Remodeling of the enhancer landscape during macrophage activation is coupled to enhancer transcription. Molecular cell, 51, 310–25.

Kall, L., Krogh, A. & Sonnhammer, E. L. 2004. A combined transmembrane topology and signal peptide prediction method. Journal of molecular biology, 338, 1027–36.

Khodor, Y. L., Menet, J. S., Tolan, M. & Rosbash, M. 2012. Cotranscriptional splicing efficiency differs dramatically between Drosophila and mouse. RNA, 18, 2174–86.

Kiontke, K., Gavin, N. P., Raynes, Y., Roehrig, C., Piano, F. & Fitch, D. H. 2004. Caenorhabditis phylogeny predicts convergence of hermaphroditism and extensive intron loss. Proc Natl Acad Sci U S A, 101, 9003–8.

Kondrashov, F. A. & Koonin, E. V. 2001. Origin of alternative splicing by tandem exon duplication. Human molecular genetics, 10, 2661–9.

Kriventseva, E. V. & Gelfand, M. S. 1999. Statistical analysis of the exon-intron structure of higher and lower eukaryote genes. Journal of biomolecular structure & dynamics, 17, 281–8.

Lalonde, E., Ha, K. C., Wang, Z., Bemmo, A., Kleinman, C. L., Kwan, T., Pastinen, T. & Majewski, J. 2011. RNA sequencing reveals the role of splicing polymorphisms in regulating human gene expression. Genome research, 21, 545–54.

Lappalainen, T., Sammeth, M., Friedlander, M. R., T Hoen, P. A., Monlong, J., Rivas, M. A., Gonzalez-Porta, M., Kurbatova, N., Griebel, T., Ferreira, P. G., Barann, M., Wieland, T., Greger, L., Van Iterson, M., Almlof, J., Ribeca, P., Pulyakhina, I., Esser, D., Giger, T., Tikhonov, A., Sultan, M., Bertier, G., Macarthur, D. G., Lek, M., Lizano, E., Buermans, H. P., Padioleau, I., Schwarzmayr, T., Karlberg, O., Ongen, H., Kilpinen, H., Beltran, S., Gut, M., Kahlem, K., Amstislavskiy, V., Stegle, O., Pirinen, M., Montgomery, S. B., Donnelly, P., McCarthy, M. I., Flicek, P., Strom, T. M., Lehrach, H., Schreiber, S., Sudbrak, R., Carracedo, A., Antonarakis, S. E., Hasler, R., Syvanen, A. C., Van Ommen, G. J., Brazma, A., Meitinger, T., Rosenstiel, P., Guigo, R., Gut, I. G., Estivill, X. & Dermitzakis, E. T. 2013. Transcriptome and genome sequencing uncovers functional variation in humans. Nature, 501, 506–11.

Lareau, L. F., Green, R. E., Bhatnagar, R. S. & Brenner, S. E. 2004. The evolving roles of alternative splicing. Curr Opin Struct Biol, 14, 273–82.

Lev-Maor, G., Sorek, R., Shomron, N. & Ast, G. 2003. The birth of an alternatively spliced exon: 3′ splice-site selection in Alu exons. Science, 300, 1288–91.

Levine, M. T., Jones, C. D., Kern, A. D., Lindfors, H. A. & Begun, D. J. 2006. Novel genes derived from noncoding DNA in Drosophila melanogaster are frequently X-linked and exhibit testis-biased expression. Proc Natl Acad Sci U S A, 103, 9935–9.

Lin, S., Coutinho-Mansfield, G., Wang, D., Pandit, S. & Fu, X. D. 2008. The splicing factor SC35 has an active role in transcriptional elongation. Nat Struct Mol Biol, 15, 819–26.

Lu, S. & Cullen, B. R. 2003. Analysis of the stimulatory effect of splicing on mRNA production and utilization in mammalian cells. RNA, 9, 618–30.

Luco, R. F., Pan, Q., Tominaga, K., Blencowe, B. J., Pereira-Smith, O. M. & Misteli, T. 2010. Regulation of alternative splicing by histone modifications. Science, 327, 996–1000.

Marques, A. C., Dupanloup, I., Vinckenbosch, N., Reymond, A. & Kaessmann, H. 2005. Emergence of young human genes after a burst of retroposition in primates. PLoS biology, 3, e357.

Mascarenhas, D., Mettler, I. J., Pierce, D. A. & Lowe, H. W. 1990. Intron-mediated enhancement of heterologous gene expression in maize. Plant molecular biology, 15, 913–20.

Matlin, A. J., Clark, F. & Smith, C. W. 2005. Understanding alternative splicing: towards a cellular code. Nat Rev Mol Cell Biol, 6, 386–98.

Merkin, J., Russell, C., Chen, P. & Burge, C. B. 2012. Evolutionary dynamics of gene and isoform regulation in Mammalian tissues. Science (New York, NY), 338, 1593–1599.

Modrek, B. & Lee, C. J. 2003. Alternative splicing in the human, mouse and rat genomes is associated with an increased frequency of exon creation and/or loss. Nature genetics, 34, 177–80.

Nielsen, C. B., Friedman, B., Birren, B., Burge, C. B. & Galagan, J. E. 2004. Patterns of intron gain and loss in fungi. PLoS biology, 2, e422.

Nott, A., Meislin, S. H. & Moore, M. J. 2003. A quantitative analysis of intron effects on mammalian gene expression. RNA, 9, 607–17.

Paten, B., Herrero, J., Beal, K., Fitzgerald, S. & Birney, E. 2008. Enredo and Pecan: genome-wide mammalian consistency-based multiple alignment with paralogs. Genome research, 18, 1814–28.

Patthy, L. 2003. Modular assembly of genes and the evolution of new functions. Genetica, 118, 217–31.

Pickrell, J. K., Marioni, J. C., Pai, A. A., Degner, J. F., Engelhardt, B. E., Nkadori, E., Veyrieras, J. B., Stephens, M., Gilad, Y. & Pritchard, J. K. 2010. Understanding mechanisms underlying human gene expression variation with RNA sequencing. Nature, 464, 768–72.

Roy, M., Kim, N., Xing, Y. & Lee, C. 2008. The effect of intron length on exon creation ratios during the evolution of mammalian genomes. RNA, 14, 2261–73.

Roy, M., Xu, Q. & Lee, C. 2005. Evidence that public database records for many cancer-associated genes reflect a splice form found in tumors and lack normal splice forms. Nucleic Acids Res, 33, 5026–33.

Roy, S. W., Fedorov, A. & Gilbert, W. 2003. Large-scale comparison of intron positions in mammalian genes shows intron loss but no gain. Proc Natl Acad Sci U S A, 100, 7158–62.

Schwartz, S., Kent, W. J., Smit, A., Zhang, Z., Baertsch, R., Hardison, R. C., Haussler, D. & Miller, W. 2003. Human-mouse alignments with BLASTZ. Genome research, 13, 103–7.

Schwartz, S., Meshorer, E. & Ast, G. 2009. Chromatin organization marks exon-intron structure. Nature structural & molecular biology, 16, 990–5.

Sela, N., Mersch, B., Gal-Mark, N., Lev-Maor, G., Hotz-Wagenblatt, A. & Ast, G. 2007. Comparative analysis of transposed element insertion within human and mouse genomes reveals Alu’s unique role in shaping the human transcriptome. Genome Biol, 8, R127.

Shabalina, S. A., Ogurtsov, A. Y., Spiridonov, A. N., Novichkov, P. S., Spiridonov, N. A. & Koonin, E. V. 2010. Distinct patterns of expression and evolution of intronless and intron-containing mammalian genes. Mol Biol Evol, 27, 1745–9.

Sharova, L. V., Sharov, A. A., Nedorezov, T., Piao, Y., Shaik, N. & Ko, M. S. 2009. Database for mRNA half-life of 19 977 genes obtained by DNA microarray analysis of pluripotent and differentiating mouse embryonic stem cells. DNA research, 16, 45–58.

Singh, J. & Padgett, R. A. 2009. Rates of in situ transcription and splicing in large human genes. Nat Struct Mol Biol, 16, 1128–33.

Sorek, R., Ast, G. & Graur, D. 2002. Alu-containing exons are alternatively spliced. Genome research, 12, 1060–7.

Sorek, R., Lev-Maor, G., Reznik, M., Dagan, T., Belinky, F., Graur, D. & Ast, G. 2004. Minimal conditions for exonization of intronic sequences: 5′ splice site formation in alu exons. Mol Cell, 14, 221–31.

Spies, N., Burge, C. B. & Bartel, D. P. 2013. 3′ UTR-isoform choice has limited influence on the stability and translational efficiency of most mRNAs in mouse fibroblasts. Genome research, 23, 2078–90.

Spies, N., Nielsen, C. B., Padgett, R. A. & Burge, C. B. 2009. Biased chromatin signatures around polyadenylation sites and exons. Molecular cell, 36, 245–54.

Sureau, A., Gattoni, R., Dooghe, Y., Stevenin, J. & Soret, J. 2001. SC35 autoregulates its expression by promoting splicing events that destabilize its mRNAs. EMBO J, 20, 1785–96.

Trapnell, C., Pachter, L. & Salzberg, S. L. 2009. TopHat: discovering splice junctions with RNA-Seq. Bioinformatics, 25, 1105–11.

Trapnell, C., Roberts, A., Goff, L., Pertea, G., Kim, D., Kelley, D. R., Pimentel, H., Salzberg, S. L., Rinn, J. L. & Pachter, L. 2012. Differential gene and transcript expression analysis of RNA-seq experiments with TopHat and Cufflinks. Nature protocols, 7, 562–78.

Wang, W., Zheng, H., Yang, S., Yu, H., Li, J., Jiang, H., Su, J., Yang, L., Zhang, J., McDermott, J., Samudrala, R., Wang, J., Yang, H., Yu, J., Kristiansen, K., Wong, G. K. & Wang, J. 2005. Origin and evolution of new exons in rodents. Genome Res, 15, 1258–64.

Wolf, Y. I., Novichkov, P. S., Karev, G. P., Koonin, E. V. & Lipman, D. J. 2009. The universal distribution of evolutionary rates of genes and distinct characteristics of eukaryotic genes of different apparent ages. Proc Natl Acad Sci U S A, 106, 7273–80.

Yeo, G. W., Van Nostrand, E., Holste, D., Poggio, T. & Burge, C. B. 2005. Identification and analysis of alternative splicing events conserved in human and mouse. Proc Natl Acad Sci U S A, 102, 2850–5.

Zhang, X. H. & Chasin, L. A. 2006. Comparison of multiple vertebrate genomes reveals the birth and evolution of human exons. Proc Natl Acad Sci U S A, 103, 13427–32.

Zhou, H. L., Hinman, M. N., Barron, V. A., Geng, C., Zhou, G., Luo, G., Siegel, R. E. & Lou, H. 2011. Hu proteins regulate alternative splicing by inducing localized histone hyperacetylation in an RNA-dependent manner. Proc Natl Acad Sci U S A, 108, E627–35.

